# *In-silico* design and assessment of OprD based multi-epitope vaccine against *Acinetobacter baumannii*

**DOI:** 10.1101/2022.05.25.493433

**Authors:** Kashaf Khalid, Saadia Andleeb

## Abstract

Gram-negative, opportunist pathogen *Acinetobacter baumannii* is notorious for causing a plethora of nosocomial infections predominantly respiratory diseases and blood-stream infections. Due to resistance development towards last-resort antibiotics, its treatment is becoming increasingly difficult. Despite numerous therapeutic developments, no vaccine is available against this ubiquitous pathogen. It is therefore apropos to formulate a rational vaccine plan to get rid of the super-bug. Considering the importance of Outer Membrane Porin D (OprD) as a potential vaccine candidate, we methodically combined the most persistent epitopes present in the *A. baumannii* strains with the help of different immunoinformatic approaches to envisage a systematic multi-epitope vaccine. The proposed vaccine contains highly immunogenic stretches of linear B-cells, cytotoxic T lymphocyte epitopes, and helper T lymphocyte epitopes of outer membrane porin OprD. The finalized epitopes proved to be significant as they are conserved in *A. baumannii* strains. The final 3D structure of the construct was projected, refined, and verified by employing several *in silico* approaches. Apt binding of the protein and adjuvant with the TLR4 suggested significantly high immunogenic potential of our designed vaccine. MD simulations showed highly stable composition of the protein. Immune simulations disclosed a prominent increase in the levels of the immune response. The proposed vaccine model is proposed to be thermostable, immunogenic, water-soluble, and non-allergenic. However, this study is purely computational and needs to be validated by follow-up wet laboratory studies to confirm the safety and immunogenicity of our multi-epitope vaccine.

## 1. Introduction

Gram-negative, opportunist *A.baumannii* is a ubiquitous coccobacillus, obligate aerobe and one of the most rampant causes of nosocomial infections typically in immunocompromised patients, accounting for enhanced mortality rates (Morris et al., 2019). Notorious for causing common outbreaks and endemics, it has been reported to transfer substantially from patient-to-patient, colonize on hands of health care workers as well as stay on environmental surfaces for longer durations, thus leading towards *A.baumannii* specific endemicity (Morris et al., 2019; Wong et al., 2016).It is one of the most common sources of myriad diseases that mainly involve bloodstream infections such as bacteremia, respiratory infections including acute pneumonia and chronic pneumonia, soft tissue infections, urinary tract infections, meningitis, and less commonly osteomyelitis (Dexter et al., 2015; Harris et al., 2019; Morris et al., 2019).

Antimicrobial and antibiotic resistance levels of *A.baumannii* are stated to be four times higher as compared to other Gram-negative bacteria such as *Pseudomonas aeruginosa* or *Klebsiella pneumoniae* (Du et al., 2019). In this context, Centers for Disease Control and Infection (CDC) has characterized the multi-drug resistant (MDR) *Acinetobacter* spp. as an urgent threat hence emphasizing the need for extensive research on therapeutics development (Giammanco et al., 2017). Additionally, *A.baumannii* has been listed by World Health Organization(WHO) among top-priority dangerous pathogens that are responsible for posing the greatest threat to public health(Harding et al., 2018).

Burgeoning resistance rates have left us with very few treatment options rendering the existing treatment approaches as mostly incompetent. Existing approaches usually involve monotherapy and combinatorial therapy which make use of different antibiotics such as polymyxins, tigecycline, and sometimes aminoglycosides. However, their undesirable pharmacokinetic properties, ability to cause toxicity (nephrotoxicity, neurotoxicity), and resistance have led to clinical failure (Shrivastava et al., 2018). Moreover, some of the antibiotics are only useful if they are used in combination whereas others are involved in increased fatality rate. Therefore, new therapeutic options are the pressing need to treat multidrug-resistant *A.baumannii* infections (Isler et al., 2018).

Taking this into account, researchers have worked extensively to devise new cost-effective therapeutic strategies that predominantly involve chemo-immuno therapies and unraveling new epitopes for active or passive immunization against the MDR pathogen. Newly proposed vaccine candidates against this pathogen include outer membrane proteins and porins like Omp34 kDa(Jahangiri et al., 2018), OprC and OmpA in combination with pal (Lei et al., 2019), outer membrane protein nuclease NucAb (Garg et al., 2016), FilF (Singh et al., 2016), BamA (Singh et al., 2017), phospholipase D (Zadeh Hosseingholi et al., 2014), Pili subunit hemagglutinin (Homenta et al., 2014) and functional exposed amino acid BauA (Sefid et al., 2015).Despite of these advancements, there have been no FDA approved vaccines in markets due to underlying limitations such as high toxicity, low immunogenicity, insolubility when expressed, or complex compositions.

Many studies have validated the use of OMPs as successful vaccine candidates in GNBs such as *Legionella pneumophila, Mannheimia haemolytica, Aeromonas salmonicida, Anaplasma marginale, Bartonella henselae, Campylobacter jejuni*, and *Avian Pathogenic E.coli (APEC)* (Ayalew et al., 2010; Diao et al., 2020; Hove et al., 2020; Moumène et al., 2015). A copious amount of information has been generated through proteomic analysis on different types of families of porins such as OmpA and OprD family in *Pseudomonas aeruginosa* (Chevalier et al., 2017). OprD is primarily involved in carbapenem uptake. Recent studies have shown downregulation of OprD and upregulation of efflux systems in the development of mutation mediated resistance (Chevalier et al., 2017; Zeng et al., 2014). The active contribution of OprD in causing the resistance makes it an important tool for combating this pathogen.

According to studies, OprD of *A.baumannii* could act as a putative vaccine candidate (Kim et al., 2016). However, there is a lack of information on the immunogenic capability of OprD in *A.baumannii*. This evidence convinces us to carry out the present study aiming at the assessment of the capacity of OprD to elicit an immune response through bioinformatics analysis. In the current study, we selected immunogenic B and T cells of OprD of *A.baumannii*. This paper concisely elucidates and explores the in-silico strategies employed in vaccine designing. Region of OprD having the greatest potential of immunogenicity has been elected as a novel candidate for the vaccine that could potentially be utilized for designing therapeutic/ prophylactic peptide vaccines against *A. baumannii*.

## 2. Materials and Methods

### 2.1 Pre-Vaccine Designing Immunoinformatic Analysis

#### 2.1.1 FASTA Sequence Retrieval

The FASTA protein sequence of OprD (ID QFQ03744) of *A.baumannii* representative strain ATCC 19606 was downloaded from National Center for Biotechnology Information (NCBI). Additional bioinformatic analysis was carried out, as described in the following section.

#### 2.1.2. Signal peptide and localization prediction

To identify possible signal peptide in the sequence, online tools SignalP 5.0 (Almagro Armenteros et al., 2019) available at http://www.cbs.dtu.dk/services/SignalP-5.0/ and LipoP (Juncker et al., 2003) at http://www.cbs.dtu.dk/services/LipoP/ were used. SignalP 5.0 encompasses artificial neural networks to improve the prediction performance of signal peptide (Almagro Armenteros et al., 2019). LipoP 1.0 locates signal peptidase I and II cleavage sites within the protein sequence (Juncker et al., 2003). Predictions about protein localization were performed by employing a template free algorithm, DeepLoc which also achieves high accuracy by employing deep neural networks (Almagro Armenteros et al., 2017). Two different servers were employed for prediction of putative transmembrane domains (TMBs): PRED-TMBB and BOCTOPUS2. PRED-TMBB enables its users to pinpoint trans-membrane strands of GNB through the method of Hidden Markov Model (HMM) (Tsirigos et al., 2016).Meanwhile, other online tool, BOCTOPUS2 predicts topology by detecting the backbone hydrogen bonding restraints that could be used to form large size TMBs (Hayat and Elofsson, 2012).

#### 2.1.3 Analysis of conserved region

A preliminary analysis of OprD protein sequence of *A.baumannii* as representative for all the possible strains of bacterium was conducted using BLASTp. BLAST search was performed with non-redundant protein sequences (nr) database of bacteria using blosum62 matrix. The retrieved sequences were aligned by multiple sequence alignment (MSA) using BLAST software, to obtain the conserved regions. To evaluate the reliability of this protein sequence, it was used as a query for performing BlastP analysis against non-redundant database restricted to Homo sapiens (taxid:9606). Non-human homologous proteins in other GNBs were also identified and interpreted to assess the inter-species and intra-species conservation of the selected protein.

#### 2.1.4. Mapping of continuous B-cell epitopes

B-lymphocytes in immune system detect and attach themselves to B-cell epitopes (BCEs) harbored within foreign molecules. Prediction of BCEs is important in designing the vaccine construct as well as diagnostic tests (Larsen et al., 2006). Several servers were used to discover potential B-Cell Epitopes. BepiPred-2.0 webserver was exploited to forecast the linear B-cells epitopes by employing random forest algorithm. As compared to other tools, this method is outstanding for sequence-based predictions (Jespersen et al., 2017). For precise estimation of the linear BCEs, Hidden Markov model, Thornton’s method, and Support Vector Machine (SVM) methods were utilized by ABCpred, BCPREDS, and SVMTriP, respectively (Saha and Raghava, 2006; Yao et al., 2020). Using various tools to find BCEs generates good quality results (Faria et al., 2011).

#### 2.1.5 Evaluation of Cytotoxic T lymphocytes (CTL)

Identifying specific peptide patterns that elicit strong MHC restricted cytotoxic T cell response is a vital step to formulate vaccine (Zhao et al., 2003). To effectively forecast the CTLs, a webserver NetCTL 1.2 was harnessed in which information is generated by artificial neural networks (ANN) and matrix methods pertaining to TAP transport efficiency, MHC class I affinity and proteasomal cleavage (Larsen et al., 2007). A default parameter 0.75 was used as a threshold value.

#### 2.1.6 Helper T-Lymphocyte (HTL) Epitope Mapping

Identification of HTLs was achieved by the webserver NetMHCII 2.3. This server calculates binding affinities of the peptides to MHC□II molecules including 7 mouse H2 alleles, 20 HLA-DQ,9 HLA-DP, and 25 HLA-DR class II alleles. It has been reported to be a highly effective tool for accurately measuring binding affinities of peptides towards MHC class II binding molecules. The results are shown in IC_50_ nM units along with the percentage rank to a set of 1,000,000 random natural peptides. IC50 binding values of < 500 nM represent strong binding affinities so this criteria was chosen (Jensen et al., 2018).

### 2.2 Multi-Epitope Subunit Vaccine Designing

Correct positioning of the nominated epitopes plays a significant role in yielding maximum immunization. Improper joining of peptides without linkers and adjuvant may lead to the synthesis of a completely new protein with unknown properties (Farhadi et al., 2015). To curb such errors, selected B-cell, HTL, and CTL epitopes were progressively connected by means of suitable linkers viz. GPGPG to connect linear B cell epitopes with HTLs and AAY to connect CTLs. Apart from enhancing the immunogenicity of the construct, these linkers are also involved in better epitope presentation and averting the production of junctional epitopes (Tahir ul Qamar et al., 2020). Adding an appropriate adjuvant is fundamental in boosting immunogenic behavior of the vaccine (Sun et al., 2018).Capability of 548AA long GroEl HSP60 of *Salmonella typhi* as an immunogenic protein has been promising (Chitradevi et al., 2013). Therefore, its protein sequence (AN: NP_458769.1) was retrieved from NCBI and saved in the FASTA format. It was attached to amino terminus of the designed construct via EAAAK linker which is a rigid helical linker that does not permit interaction of construct with other areas of proteins, thus resulting in stability (Choi et al., 2019).

### 2.3. Post-Design Analysis and Validation

#### 2.3.1. Assessment of Physicochemical Aspects and Peptide Solubility

The physicochemical aspects of designed vaccine construct was predicted by the ExPasy Protparam webserver. The parameters calculated were the total number of residues, molecular weight in kDa, in-vivo and in-vitro half-life, aliphatic index, theoretical pI, instability index, and grand average of hydropathicity (GRAVY) (Gasteiger et al., 2005). To gauge the solubility of the peptide, Protein-Sol webserver was employed. This webserver uses a fast protein sequence-based calculation of solubility. The prediction output is given in the format of 0-1 range with values greater than the average value of 0.45 being considered as soluble (Hebditch et al., 2017).

#### 2.3.2. Immunogenicity Profiling

Allertop v 2.2 was employed to investigate the allergenicity of the vaccine. It uses a machine learning method that classifies protein using the k-nearest neighbors (kNN) algorithm and thus shows accuracy level of 85.3% at 5-fold cross-validation (Dimitrov et al., 2014). VaxiJen v 2.0 and ANTIGENpro tools to measure antigenic behavior of the peptide were utilized. VaxiJen depicts the immunogenic potential of construct with an accuracy ranging from 70% to 89% (Flower et al., 2017). According to several reports, ANTIGENpro can predict the protein antigenicity with an accuracy greater than 75.5% (Magnan et al., 2010).

#### 2.3.3. Determination of the Secondary Structure

To determine the secondary structure of the polypeptide, a webserver PSIPRED was employed which performs processing of position specific scoring matrix using two feed forward neural network with an accuracy of 84.2% (McGuffin et al., 2000).

#### 2.3.4 Derivation of 3-Dimensional Structure of Vaccine

To forecast the 3D shape of the peptide, webserver I-TASSER (Iterative Threading ASSEmbly Refinement) was utilized that performs the structure folding and remodeling through Monte Carlo simulations which are based on the improved knowledge-based force field. It consists of three key methodologies: (i) hydrogen-bonding networks (ii) basic statistical potentials, and (iii) threading-based restraints from LOMETS (Yang et al., 2014). It is reported to be a top-ranked server for carrying out structural predictions for the last five CASP experiments (Zhang, 2008).

#### 2.3.5. Improvement of Derived Protein Structure

The initial models are further sent for improvement that includes accurate and quality transformation of initial models into structures that are comparable with experimental structures (Feig, 2017). ModRefiner server http://zhanglab.ccmb.med.umich.edu/ModRefiner was utilized for preliminary refinements to achieve an overall improvement in the physical structure (Xu and Zhang, 2011). Next, a webserver GalaxyRefine http://galaxy.seoklab.org/cgi-bin/submit.cgi?type=REFINE was employed to further enhance the structural quality. This method involves remaking and repacking of side chains as well as overall structure relaxation by molecular dynamics simulations. It has refinement (Heo et al., 2013).

#### 2.3.6 Structure Validation

Evaluation and confirmation of the structural quality of the protein construct are ensured via several quality assessment tools such as Ramachandran plot analysis for viewing torsion angles distribution in the structure, clash scores for all-atom contact analysis, aberrations in structural conformity, and rotamers analysis. These indicators have proved to be highly valuable in determining the accuracy of local and global three-dimensional structures of protein (Pražnikar et al., 2019). In light of this, many online web servers were utilized to confirm the refined structures. These include (i) ProSA-web https://prosa.services.came.sbg.ac.at/ (Wiederstein and Sippl, 2007) (ii) ERRAT https://servicesn.mbi.ucla.edu/ERRAT/ (Dym et al., 2012) (iii) MolProbity http://molprobity.biochem.duke.edu/ (Chen et al., 2010). PROCHECK https://servicesn.mbi.ucla.edu/PROCHECK/ (Roman Laskowski et al., 1983) was also employed to analyze the Ramachandran Plot. The Ramachandran plots help in analyzing the dispersal of (φ, Ψ) torsion angles of the protein backbone to evaluate the protein structures (Sobolev et al., 2020).

#### 2.3.7 Mapping of discontinuous B-cell epitopes (Conformational)

Distantly located residues on the primary structure of a protein that fall into proximity due to folding of a protein structure constitute >90% of discontinuous B-cell epitopes (Mukonyora, 2015). An online tool, ElliPro http://tools.iedb.org/ellipro/ was employed to infer conformational BCEs. Ellipro is considered the state of the art webserver as it gave an AUC value of 0.732 (Ponomarenko et al., 2008).

#### 2.3.8 Molecular Dynamics (MD) Analysis

GROMACS 5.0 software was used to have a deep understanding of structural integrity of protein in a life-like simulated environment (Abraham et al., 2015). OPLS-AA force field was applied to the system and overall charge was noted when pdb file was used as an input. SPC/E water model was used to solvate the cubic system and genion tool was employed to add ions into the system to nullify the overall charge. It is pertinent to stabilize the temperature using NVT ensemble and pressure under NPT ensemble. Subsequently, MD simulations were performed for 10 nanoseconds (ns) and resulting trajectories of root-mean square deviation (RMSD) and root-mean square fluctuations (RMSF) were examined.

#### 2.3.9. Molecular Docking Analysis

Only appropriate attachment of immunogenic molecules to the specific immune receptors can evoke a proper immune response. TLR4 is involved not only in generating suitable immune response and long-lasting immunity but also plays a specific role in *A. baumannii* infections *in vitro* and *in vivo* (Monem et al., 2020; Pulido et al., 2020; Shi et al., 2020). Therefore, TLR4 was selected as an immune receptor for binding to the vaccine construct. TLR4 binding pockets were forecasted by the CASTp v 3.0 server (Computed Atlas of Surface Topography of proteins) accessible at http://sts.bioe.uic.edu/castp/. Docking analysis of the designed construct with TLR4 (PDB ID:4G8A), was conducted by a python-based web server HADDOCK v 2.4 (High Ambiguity Driven protein-protein DOCKing) available at https://wenmr.science.uu.nl/haddock2.4/. Interactions are measured in the terms of inter and intramolecular energies that involve Van der Waal forces and electrostatic energies (Ambrosetti et al., 2020). Three steps are involved in the docking protocol viz, stringent body energy minimization, an intermediately-flexible fine-tuning in torsion angle space, and an ultimate refinement in the explicit solvent (De Vries et al., 2010). To accurately drive the docking protocol, active and passive residues were procured using the online web-tool CPORT (Consensus Prediction Of interface Residues in Transient complexes) https://milou.science.uu.nl/services/CPORT/ (Xue et al., 2016). Thus, the data collected after successfully predicting the active and passive residues was used to guide the prediction-driven docking in HADDOCK v 2.4 to better explore the interactions among the vaccine construct and TLR4 as well as the adjuvant and TLR4. The structure of adjuvant GroEl (AN: NP_458769.1) was modeled by employing SWISS-Model (https://swissmodel.expasy.org/). To calculate the protein binding affinities, PRODIGY web-server (PROtein binDIng enerGY prediction) (Xue et al., 2016) accessible at https://bianca.science.uu.nl/prodigy/ was exploited.

#### 2.3.10. Codon analysis and mRNA expression of the target protein

Codon optimization is essential to promote copious yields of gene product in the host. Codon adaptation as well as the reverse translation was performed to improve the gene expression levels in the host. Therefore, codon utilization of the intended construct in *Escherichia coli* K12 strain was performed by the JCat (Java Codon adaptation tool) server at http://www.jcat.de/ (Grote et al., 2005). Moreover, three additional opportunities were availed to avoid cleavage sites of restriction enzymes, rho-independent transcription terminators, and prokaryotic ribosome binding. The results show enhanced GC content besides Codon Adaptation Index (CAI), which indicates expression levels of protein. CAI score of 1 is considered ideal value however values greater than 0.8 are deemed desirable. To enhance efficacy and promote expression of the designed peptide in host, it was cloned in *E.coli* vector by integrating two restriction sites viz., *Nde I* and *Xho I* at the amino and carboxy-terminus of the designed construct, respectively. Finally, the adjusted sequence was incorporated into the pET30a (+) expression vector via the SnapGene software.

#### 2.3.11. Estimation of the immune response

A webserver C-ImmSim available at http://150.146.2.1/C-IMMSIM/index.php generates simulations of the responses generated by the immune system upon stimulation by the final vaccine construct. It produces simulations for both humoral and cellular immune responses that are markedly tantamount to the actual immune responses generated by the human body (Rapin et al., 2010). Three simulated injections having no LPS were given at three time intervals 1,42 and 84 using default settings of simulation volume and simulation steps set at 10 and 100 respectively and the random seed at 12345 (Majid and Andleeb, 2019).

## 3. Results

### 3.1. Pre-Vaccine Design Analysis

#### 3.1.1. Sequence retrieval and primary *in-silico* analysis

The complete protein sequence of OprD (AN: QFQ03744) 438 amino acid in length, belonging to representative strain ATCC 19606 of *A.baumannii* was downloaded in FASTA format from NCBI. The protein was predicted to be extracellular by Deeploc. Signal peptide was identified at the position 1 to 22 of the sequence with position 22-23 as a cleavage site which could be cleaved by signal peptidase. BOCTOPUS2 and PRED-TMBB predicted eight transmembrane strands. BLASTp analysis was carried out using the accession number as a query sequence against which no hit specific to the taxid:9606 Homo sapiens was found. However, the protein sequence covered 726 strains of *Acinetobacter* spp and thus suggested wide-spread presence of this protein sequence amongst numerous strains of *A.baumannii*. Results revealed 93.76% to 100% similarity among *A.baumannii* strains by covering 726 strains of *A.baumannii* and found 40 hits of OprD which shared ≥97% identity, 100% query coverage, and E-value of 0, thus proving as a universally conserved *A. baumannii* antigen. OprD is also found in *Pseudomonas aeruginosa* and *Klebsiella pneumoniae* strains but with lesser identity (36-40% and 37% respectively).

#### 3.1.2. Prediction of linear B-cells

The selection of the most recurrent and immunogenic epitopes was done carefully to design the vaccine construct. BCPred, ABCpred, SVMTrip and BepiPred 2.0 predicted a total of 74 epitopes. and final recruitment of selected epitopes in the development of a new vaccine construct (Table 1).

**Table 1.**
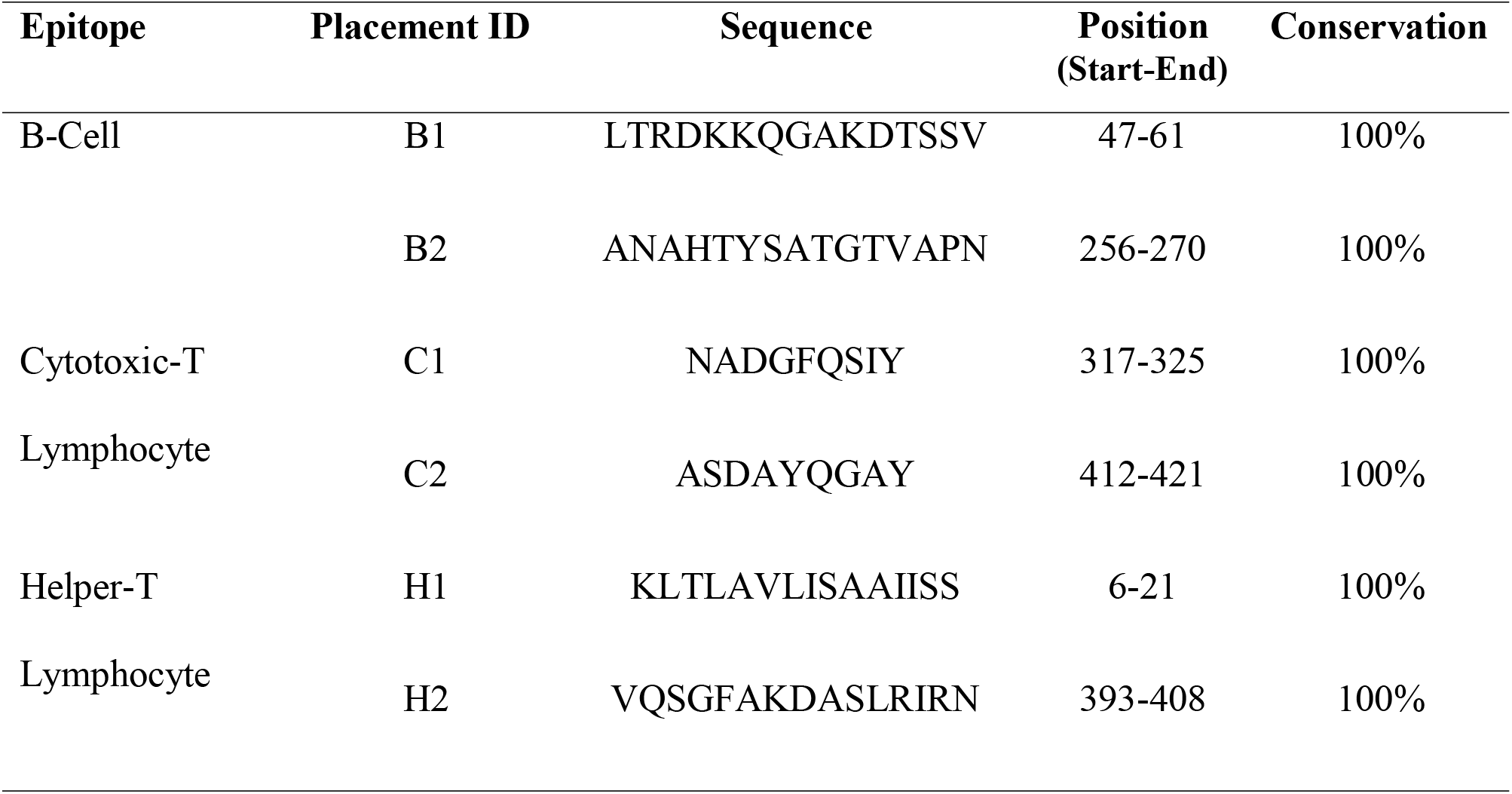
The final selected epitopes, their position, and Placement ID in the vaccine design.

#### 3.1.3. Prediction of Cytotoxic T Lymphocytes (CTLs)

Overall, 20 CTL (9-mer) epitopes were identified with the assistance of NetCTL 1.2 webserver. Default settings were selected for the calculation of epitopes. Finally, 02 epitopes were carefully chosen based on their high-affinity scores (Table 1).

#### 3.1.4. Prediction of Helper T Lymphocytes (HTLs)

The webserver NetMHCII 2.2 envisaged MHC-II epitopes that illustrate high binding scores towards the HLA alleles in the form of IC50 values (in nM). For final incorporation into the vaccine construct, a sum of two epitopes were opted (Table 1).

### 3.2 Construction of final vaccine with different adjuvants and linkers

To design an efficient vaccine, it is necessary to make use of a potent adjuvant that would result in a boosted immunogenicity potential of the construct. A total of 06 peptides were arranged and the GroEl adjuvant from Salmonella typhimurium was incorporated into the amino terminus of the B-cell epitopes using the EAAAK linker. MHC-1, MHC-II epitopes, and B cell epitopes were connected via specific linkers, AAY, and GPGPG as they do not modify the conformation of vaccine construct. EAAAK linker helps in the attachment of the adjuvant at the N-terminus of B-cell epitope. The remaining BCEs and HTLs were joined together with the help of GPGPG linkers. For proper fitting of the CTL epitopes, AAY linkers were utilized. At the carboxy terminus of peptide, six-His tag was attached to promote the process of identification and purification (Fig. 2)

**Figure 1.**
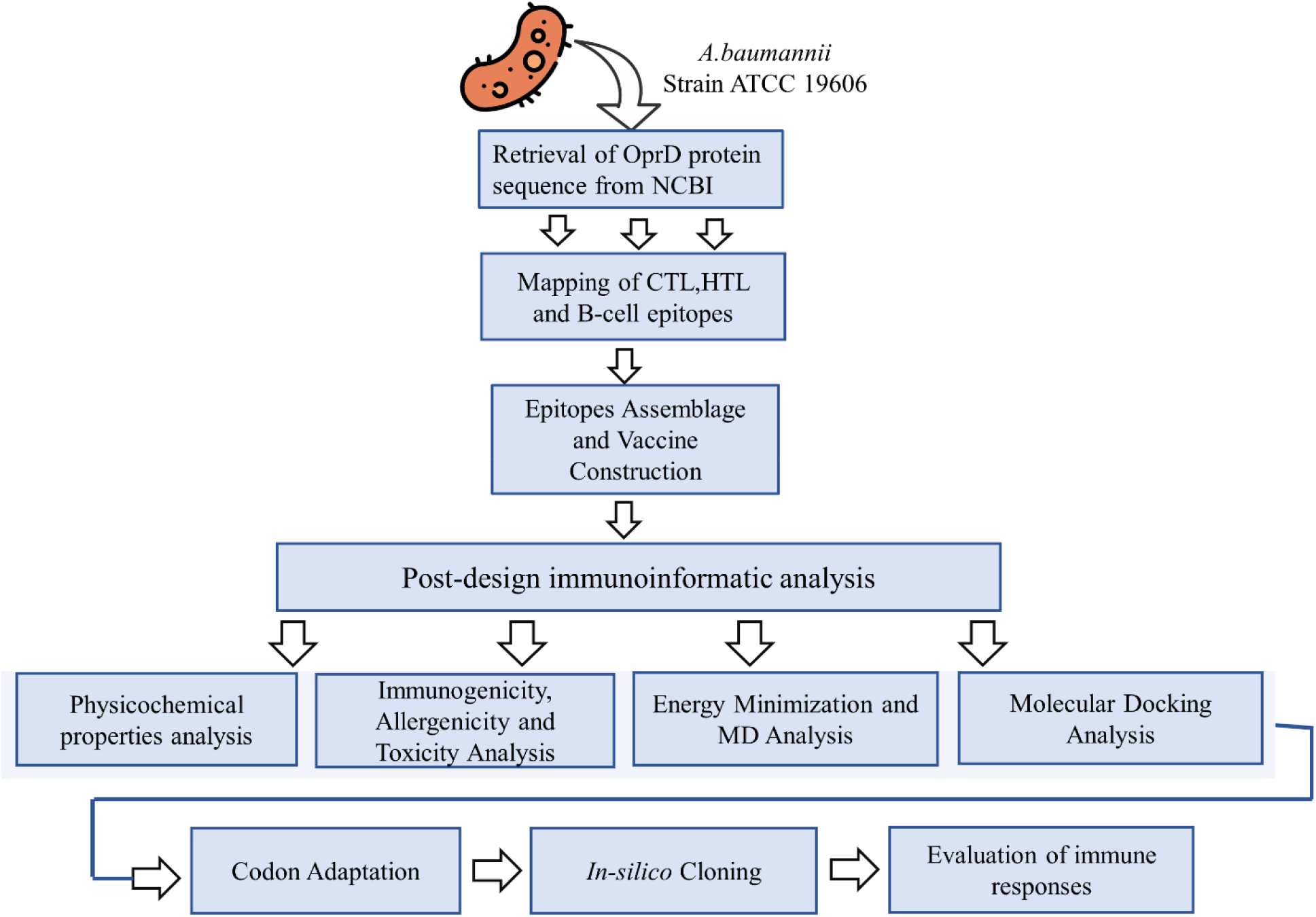
General schema of the strategy followed in the present study.

**Figure 2.**
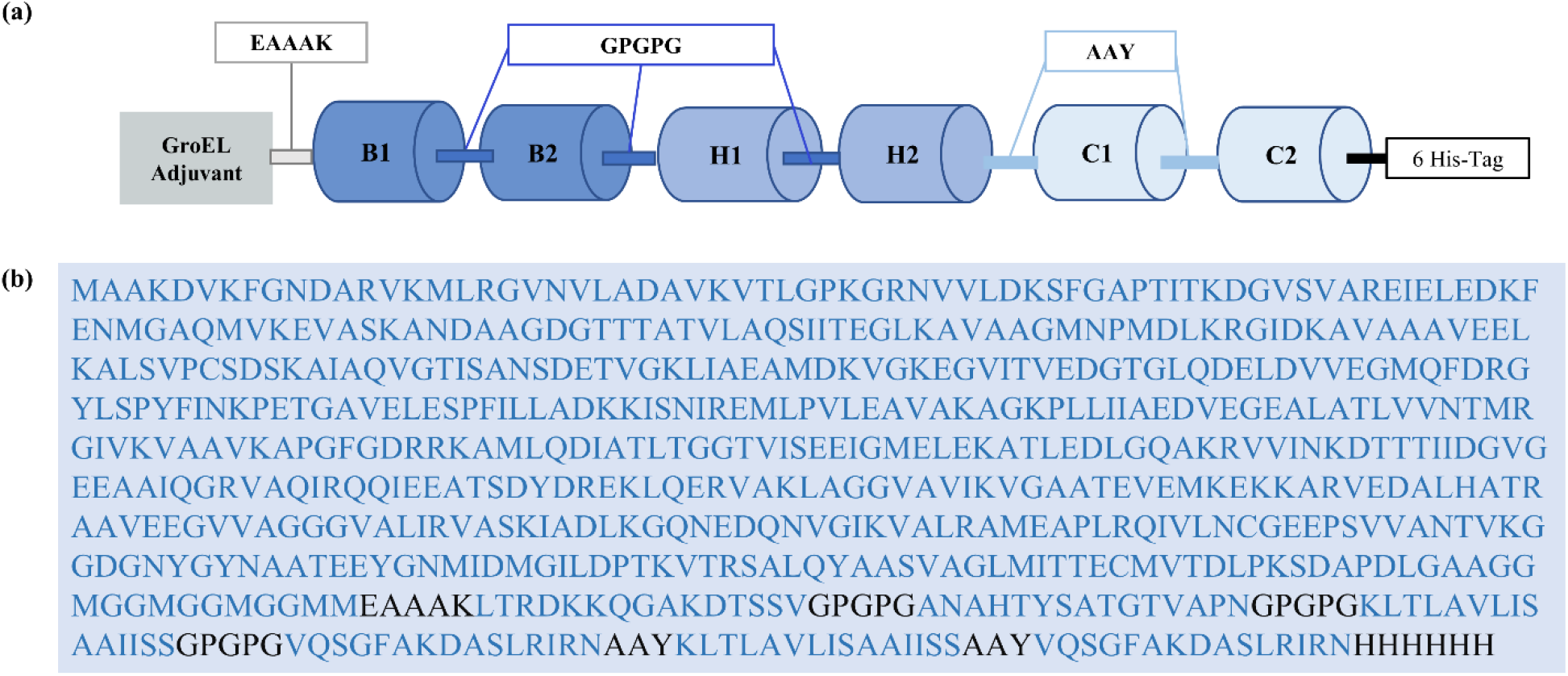
Schematic diagram and amino acid sequence of the final vaccine construct. (a) Arrangement of selected epitopes along with adjuvant and appropriate linkers. (b) Blue ink represents the amino acid sequences of epitopes and adjuvant whereas black ink represents the linkers and 6-His tag.

### 3.3 Post-Vaccine Design Analysis and Validation

#### 3.3.1. Analysis of physicochemical features

The physicochemical aspects were estimated using the FASTA format in the ExPASY ProtParam webserver to know about the basic characteristics of the vaccine. The molecular weight was calculated to be 69.9 kDa. The isoelectric point value (pI) was found to be 5.51, which illustrates that the protein is slightly acidic. ProtParam declared the protein as stable by identifying the instability index (II) value as 28.26.II values greater than 40 signify instability. The GRAVY value was noted to be −0.026 exhibiting the hydrophilic nature of the protein and thus the possibility of interactions with the neighboring water molecules. Half-life of the designed construct was evaluated to be 30 hours in mammalian reticulocytes in vitro, >20 hours in yeast, and >10 hours in E. coli in vivo. The protein displays the property of being soluble when expressed by exhibiting the solubility score of 0.611. Furthermore, the protein construct was indicated to be thermostable by showing the aliphatic index value of 74.85.

#### 3.3.2. Antigenicity, Toxicity, Allergenicity

The overall antigenicity of the construct was estimated to be 0.64 in bacteria and 0.55 in virus model by VaxiJen. AntigenPro estimated the antigenicity score to be 0.88. The VaxiJen score at a threshold value of 0.4 was also determined excluding the adjuvant for which the antigenicity score was estimated 0.78 in the bacteria model and 0.477 in the virus model. It is clear from the results that the vaccine construct itself can elicit the immune response whether an adjuvant is attached. The protein was proposed to be a non-allergen by AllerToP v 2.0.

#### 3.3.3. Secondary Structure

The final construct consisted of 670 residues with 34.48% of the random coil, 54.03 % of the alpha-helical structure, and 11.49 % of the extended strand (Fig. 3)

**Figure 3.**
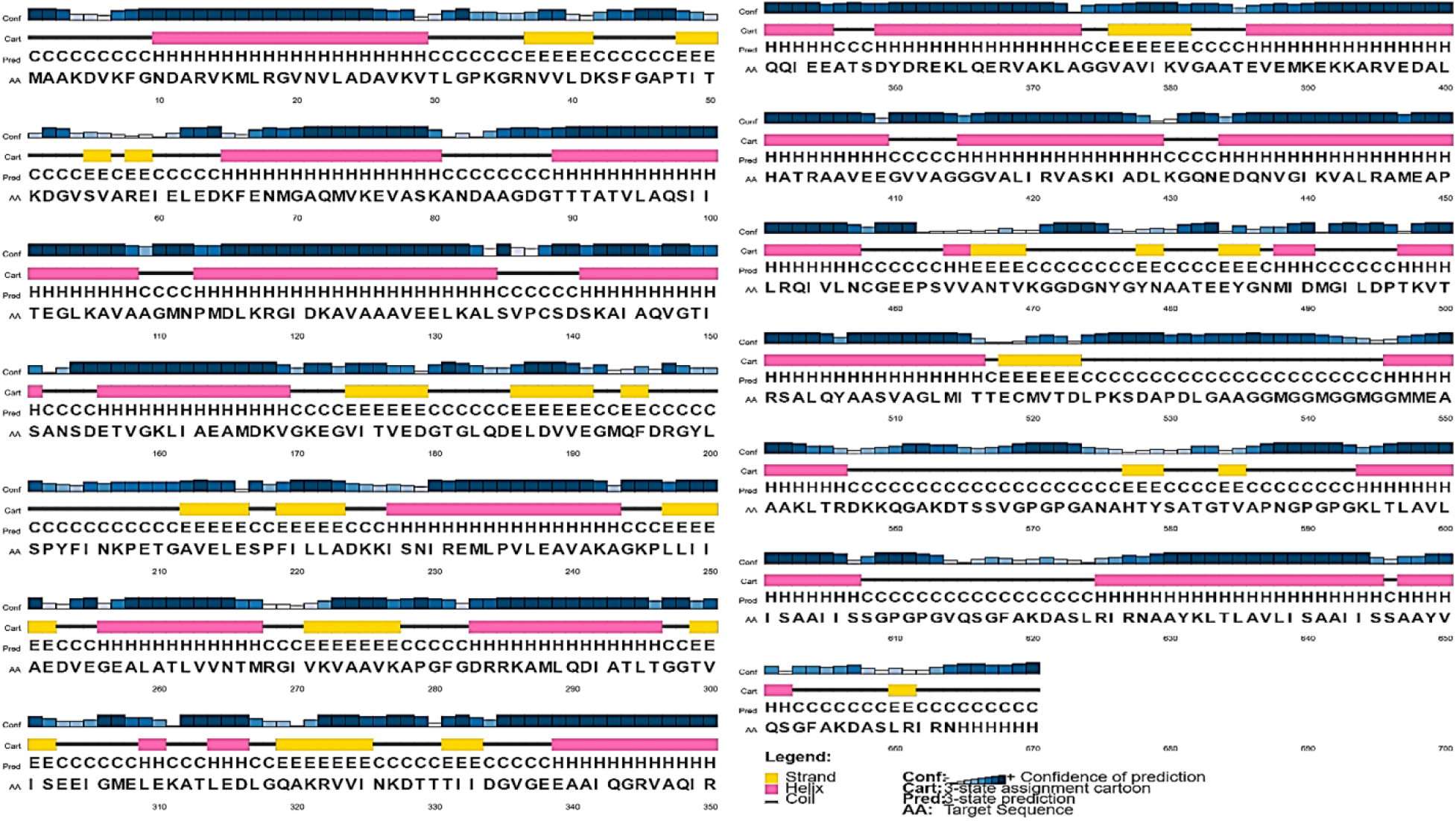
PSIPRED generated secondary structure of the designed peptide. (a) E denotes extended strand, h: alpha helix, c: random coil, and b: Beta turn. (b) Elements of secondary structure (blue: alpha helix, red: extended strand, green: beta turn, yellow: random coil

#### 3.3.4. Derivation of the Tertiary Structure

I-TASSER server predicted five models based on top ten threading templates which exhibited good alignment according to their z-scores ranging from 1.71-15.61. The predicted models showed c-scores in the range of -2.50 to -0.45. The normal value range of the C-score is -5 to 2, where values closer to 2 represent higher confidence in the structure. The predicted structure showing the maximum c-score of -0.45 was chosen for additional refinements (Fig 4a). The TM score and RMSD value were documented to be 0.66 and 3.0 ± 4.6, respectively. A model demonstrates precise topology if the TM value is greater than 0.5. Values lesser than 0.17 indicate nonspecific similarity.

**Figure 4.**
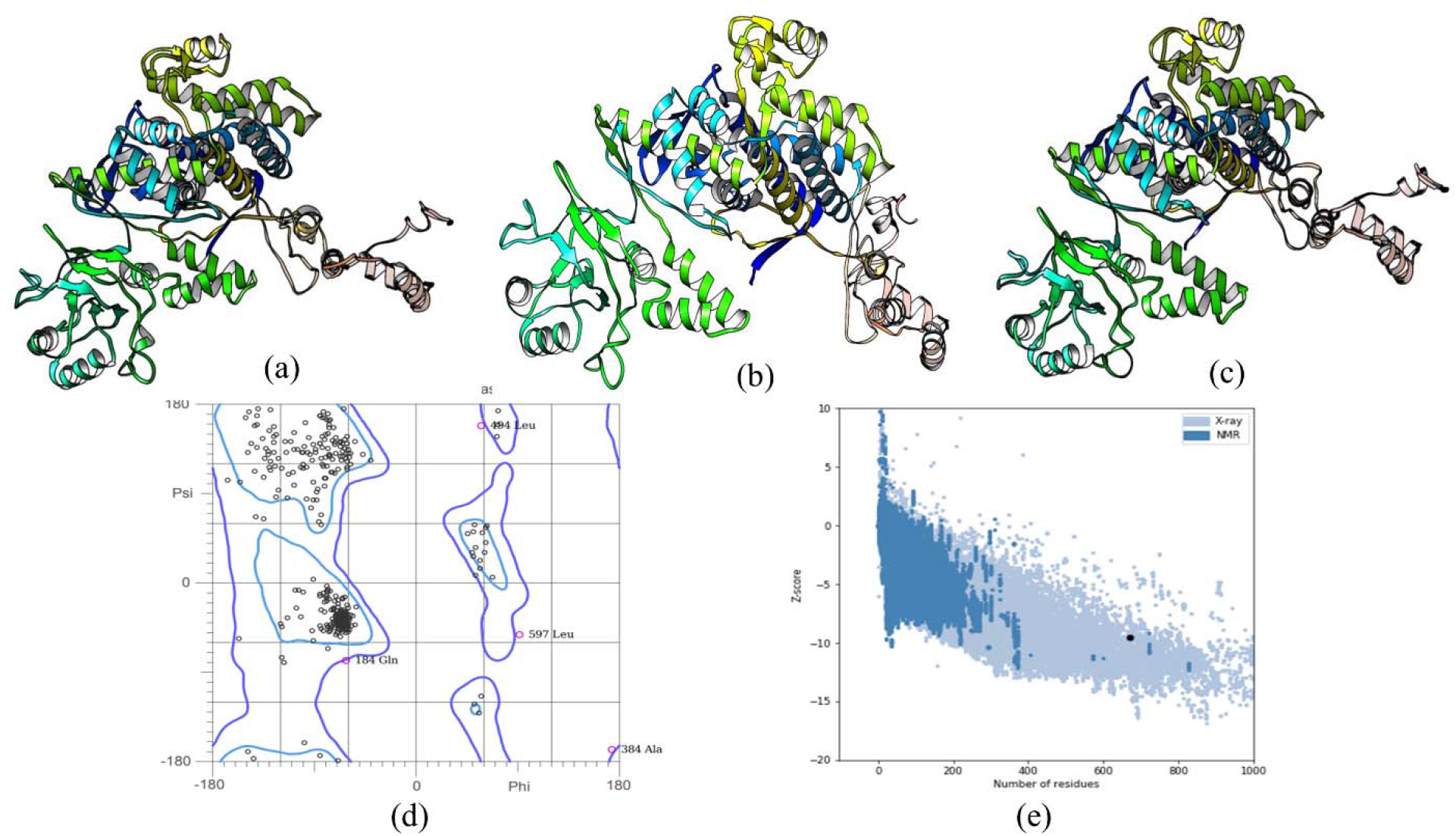
3-D structure prediction, improvement, and validation. (a) 3-D structure modeled by I-TASSER (b) structure refined by ModRefiner (c) Improved structure by GalaxyRefine (d) Ramachandran Plot depicting allowed and disallowed regions (e) Z-score scatter plot

#### 3.3.5. Refinement of Protein 3-D Structure

Rectification of the primary vaccine model was attained by the ModRefiner server (Fig 4b) followed by subsequent refinements by GalaxyRefine (Fig. 4c). After several refinements, a final model was selected based on distinct parameters which include RMSD (0.286), GDT-HA (0.9873) and MolProbity (2.046). Poor rotamers value and clash score was observed to be 0.0 and 10.7, respectively. Ramachandran plot score was noted to be 94.5%. Therefore, further immunoinformatic analysis were performed on this carefully chosen model.

#### 3.3.6. Structure Validation

According to the Ramachandran plot analysis, 94.46 % of protein residues were found to be present in the favored region (Fig. 4d). This value is consistent with the GalaxyRefine score of 94.5%. Moreover, 8.7% residues were observed in the allowed region and only 1% of the residues were detected in the disallowed region. These percentages symbolize the good quality of our predicted model. To determine the presence of any possible errors in the refined model or validate the global quality of the structures and, it is necessary to determine ERRAT and ProSA-web scores. Thus, the Z-score of -9.54 (Fig. 4e) and the quality factor of 87.6% further endorse the good quality of our model.

#### 3.3.7. Predicted Discontinuous Epitopes

A sum of 7 discontinuous B-cell epitopes having 127 residues in total and the immunogenicity scores ranging from 0.641 to 0.891 was identified. Varying sizes of the conformation epitopes were documented extending from five to thirty-nine residues (Table 2, Figure 5).

**Table 2.** Discontinuous epitopes present in the protein as predicted by the ElliPro. A total of 127 residues lie in the 7 conformational B-cell epitopes

| No. | Residues | No. of residues | Score |
| --- | --- | --- | --- |
| 1 | A:L633, A:T634, A:L635, A:A636, A:V637, A:L638, A:I639, A:S640, A:A641, A:A642, A:I643, A:I644, A:S645, A:S646, A:A647, A:A648, A:Y649, A:V650, A:Q651, A:S652, A:G653, A:F654, A:A655, A:K656, A:D657, A:A658, A:S659, A:L660, A:R661, A:I662, A:R663, A:N664, A:H667, A:H669, A:H670 | 35 | 0.966 |
| 2 | A:H665, A:H666, A:H668 | 3 | 0.965 |
| 3 | A:A619, A:K620, A:D621, A:A622, A:S623, A:L624, A:R625, A:I626, A:R627, A:N628, A:A629, A:A630, A:Y631, A:K632 | 14 | 0.92 |
| 4 | A:A550, A:A551, A:A552, A:K553, A:L554, A:T555, A:R556, A:D557, A:K558, A:K559, A:Q560, A:G561, A:A562, A:K563, A:D564, A:T565, A:S566, A:S567, A:V568, A:G569, A:P570, A:G571, A:P572, A:G573, A:A574, A:N575, A:A576, A:H577, A:T578, A:Y579, A:S580, A:A581, A:T582, A:G583, A:T584, A:V585, A:A586, A:P587, A:N588, A:G589, A:P590, A:G591, A:P592, A:G593, A:K594, A:L595, A:T596, A:L597, A:A598, A:V599, A:L600, A:I601, A:S602, A:A603, A:A604, A:I605, A:I606, A:S607, A:S608, A:G609, A:P610, A:G611, A:P612, A:G613, A:V614, A:Q615, A:S616, A:G617, A:F618 | 69 | 0.775 |
| 5 | A:G198, A:Y199, A:L200, A:S201, A:P202, A:Y203, A:F204, A:I205, A:N206, A:K207, A:P208, A:E209, A:T210, A:G211, A:A212, A:V213, A:E214, A:L215, A:E216, A:S217, A:P218, A:F219, A:L221, A:A223, A:D224, A:K225, A:K226, A:I227, A:S228, A:N229, A:I230, A:R231, A:E232, A:M233, A:L234, A:P235, A:V236, A:L237, A:E238, A:A239, A:V240, A:A241, A:K242, A:A243, A:G244, A:K245, A:P246, A:L247, A:L248, A:I250, A:A251, A:E252, A:D253, A:V254, A:E255, A:G256, A:E257, A:A258, A:L259, A:A260, A:T261, A:L262, A:V263, A:V264, A:N265, A:T266, A:M267, A:R268, A:G269, A:I270, A:V271, A:K272, A:V273, A:A274, A:A275, A:V276, A:K277, A:A278, A:P279, A:G280, A:G282, A:D283, A:R284, A:G298, A:T299, A:V300, A:I301, A:S302, A:E303, A:E304, A:I305, A:G306, A:M307, A:E308, A:L309, A:E310, A:K311, A:A312, A:T313, A:L314, A:E315, A:D316, A:L317, A:G318, A:Q319, A:A320, A:K321, A:I325, A:N326, A:K327, A:D328, A:T329 | 112 | 0.684 |

**Figure 5.**
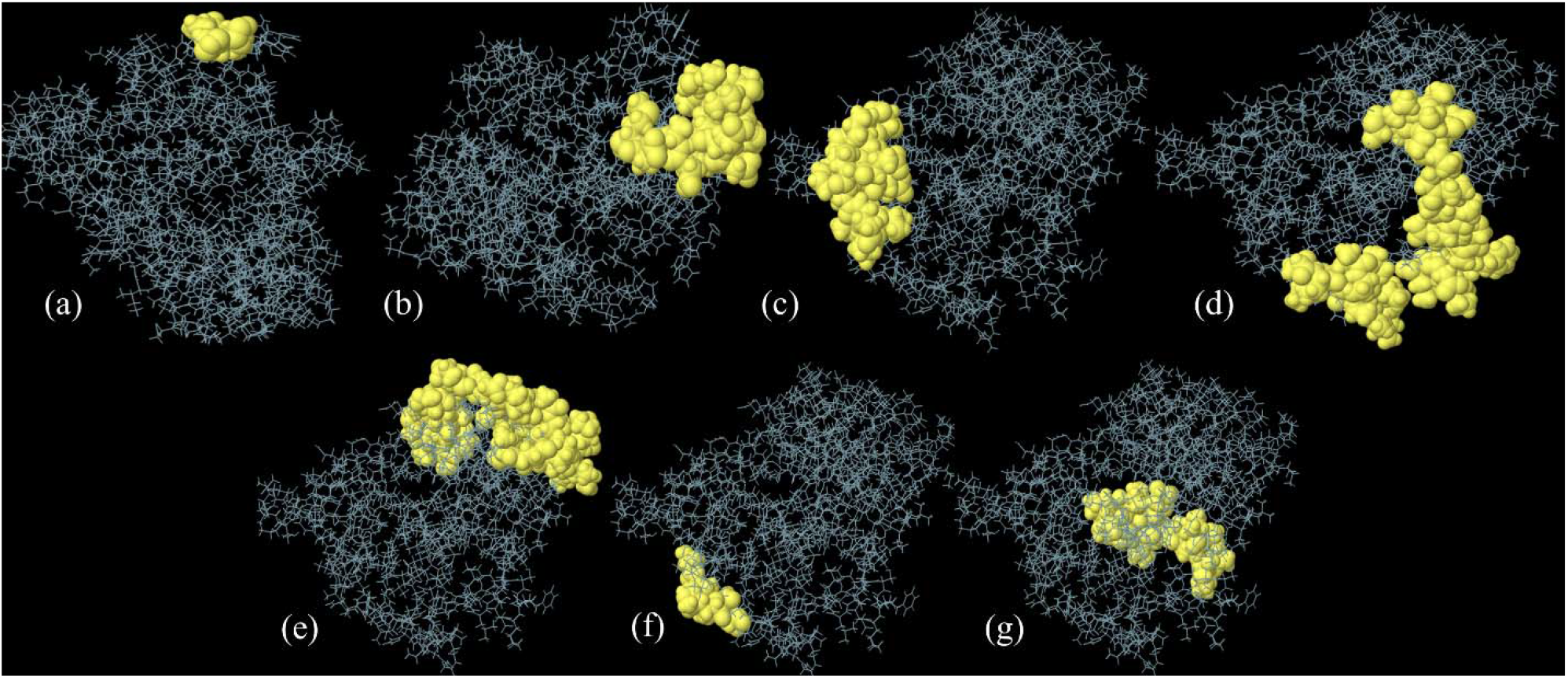
Conformational B-cell epitopes projected as by ElliPro. (a-g) Depiction of conformational B-cell epitopes of the OprD protein of *A. baumannii* from different angles. The yellow portion shows the epitopes, rest of the protein is symbolized by grey sticks.

#### 3.3.8. Molecular dynamics analysis of multi-epitope vaccine

To assess the movement of atoms and stability of protein, molecular dynamics simulation was performed using the GROMACS server. The designed system underwent equilibrations (potential energy, pressure, and temperature) followed by energy minimization. As a result, graphs revealed that system reached a desirable temperature of 299.7 K (Fig. 6a) and an average pressure of 0.92 bar with a total drift value of 0.62(Fig. 6b). To further analyze the overall stability of the structure and how much the conformation was changed between two time points, RMSD graph was analyzed. It showed that the protein remained highly stable. Initially the fluctuations started from 0.12 nm and go up to 1.10 nm in time of 8.8 ns. These slight oscillations show that model maintains stability over the period (Fig. 6c). To further analyze the conformational stability, RMSF plot was generated and analyzed in which the highest peak of fluctuation was observed to be at 1.37 nm, while the lowest peak of fluctuation was at 0.15 nm. The overall graph showed very slight oscillations where higher peaks in graph depict regions of higher flexibility (Fig. 6d).

**Figure 6.**
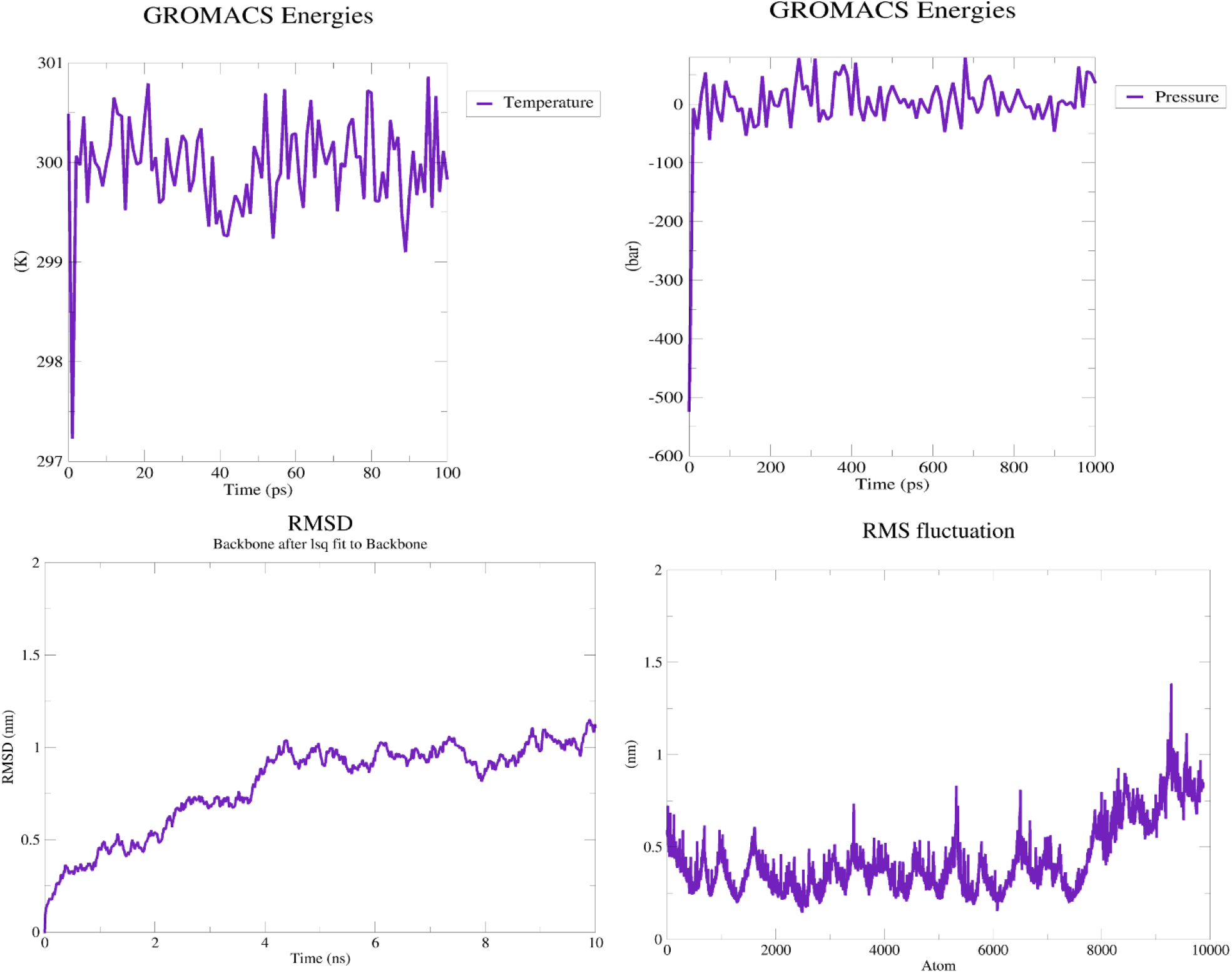
Molecular dynamics simulation of vaccine (a) Temperature equilibration (at 100 picoseconds) of the simulated environment obtained via classical NVT ensemble shows average value of 300 (K) (b) Pressure equilibration (at 100 picoseconds) of the simulated environment obtained via classical NPT ensemble shows average pressure of 0.92 bar (c) RMSD plot of the vaccine shows slight oscillations, confirming the overall protein stability (d) RMSF plot revealing flexibility of the protein through high fluctuation peaks.

#### 3.3.9 Molecular Docking

HADDOCK v 2.2 was used harnessed to perform the molecular docking using the data-driven approach. To understand the contacts between the immune receptor and the TLR4, the binding pockets of the protein were estimated by the CASTp 3.0 server. As a result, a binding pocket with a molecular surface area of 2933.3 Å2 and a molecular surface volume of 7317.6 Å3 was identified. This pocket having the mouth molecular surface of 418.2 Å2 and the molecular surface circumference sum of 348.1 Å could act as a potential interacting surface. In order to gain insights into the residues actively participating in the interaction, CPORT gave a list of active residues A2,K3,L4,T6,E8,V39,A43,A44,G45,A46,A108,S200,A202,A203,I204,S205,A256,S257,I260,R 261,H263,H264 from the B chain of the GroEL adjuvant, T6, E8, E29, V33, A36, A37, V39, A43, A44, G45, A46, A47, A49, K108, E109, D112, A114, E199, K277, F279, G280, D281, M285, Q288, T329, I330, D332, Y476, E481, N485,M486, M489, G490, D493, I513, E516, C517, M518, V519 from the A chain of the chimeric protein and E26, E41, L42, N43, Y45, K46, I47, P48, N50, L51, F53, S54, T55, H67, L68,S70, Y71, S72, F73, S75, F76, E78, L79, S99, S100, S103, S124, Y189, T191, L305,S384, G408, V409, V440, K475, A477, D500, S502, N524, S526, L533, D534, D548, S550, M555,S557, N573, T575, A580 from the B chain of TLR4. These active residues were fed to server to drive docking protocol smoothly and effectively. The HADDOCK results generated several docked complexes out of which the clusters having the lowest RMSD values (protein-TLR4 complex (0.5 Å), adjuvant-TLR4 complex (11.6 Å) with the best poses and least intermolecular energies among the construct-TLR4 complex (−317.5 Kcal/mol), adjuvant-TLR4 complex (−389.2 Kcal/mol) were carefully chosen. According to the results generated by the PRODIGY, the relative binding free energies (ΔG) of both complexes; construct-TLR4 (−16.7Kcal/mol) and adjuvant-TLR4(−11.8Kcal/mol) exhibit the capacity of construct and adjuvant to bind to each other properly and thus playing role in instigating the TLR4 receptor response by making suitable conformational changes. Additionally, a total of 11 hydrogen bonds between the vaccine-TLR4 complex as well as 14 hydrogen bonds between the adjuvant-TLR4 complex indicate strong bonding. The examination of the number of interfacial contacts (IC) per property between the two complexes was (ICs charged-charged: 13, ICs polar-polar: 14, ICs apolar-apolar:23) in case of vaccine and TLR4 complex whereas for the adjuvant and TLR4 complex values are (ICs charged-charged: 14, ICs apolar-apolar: 20, ICs polar-polar:1) (Figure 7, Figure 8).

**Figure 7.**
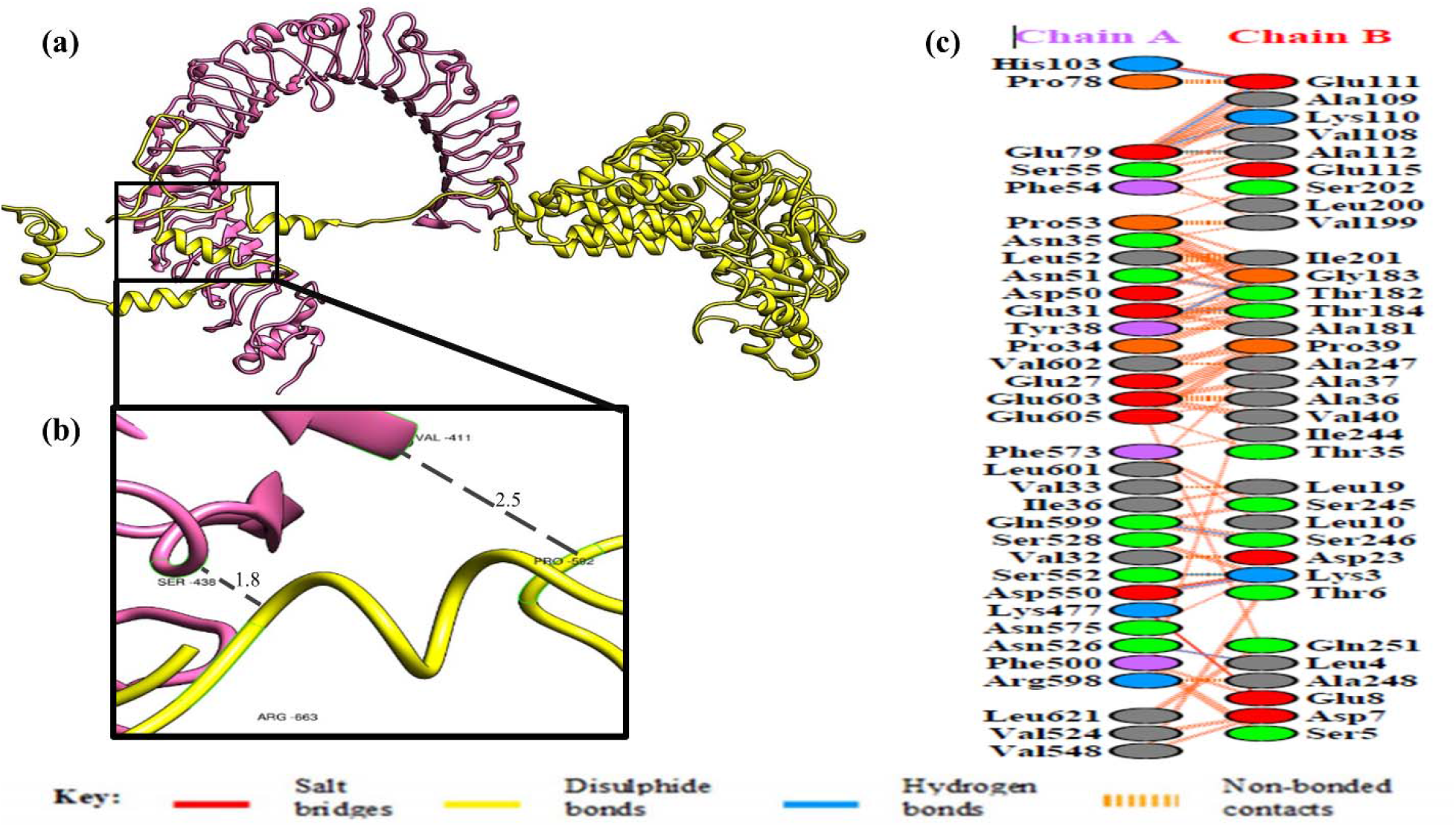
Molecular docking of the designed construct with TLR-4. (a) Binding of TLR-4 (Hot-pink) with vaccine model (Yellow). (b)The interface of binding residues. (c) Interacting bonds among the vaccine and TLR-4

**Figure 8.**
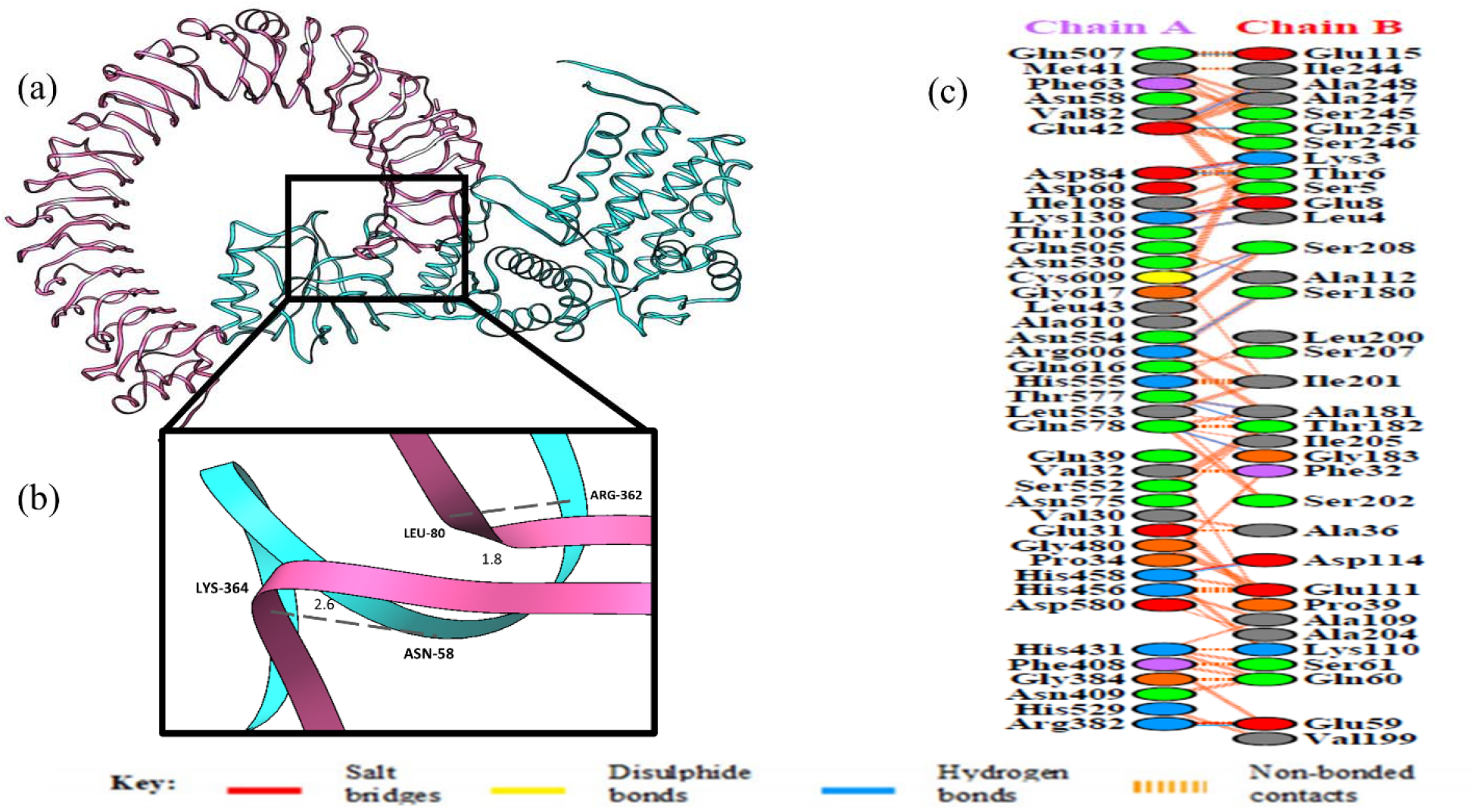
Molecular docking of GroEl adjuvant with TLR-4. (a) Binding of TLR-4 (Hot-pink) with GroEl adjuvant (Cyan). (b)The interface of binding residues. (c) Interacting bonds among the vaccine and TLR-4

#### 3.3.10. cDNA and mRNA Analysis for Cloning and Expression Prediction

The frequency of codon usage of the recombinant protein as well as the expression host needs to be similar. For this purpose, codon optimization of the designed construct is performed utilizing the JCat (Java Codon Adaptation Tool). The improved sequence was 2033 nucleotides long having a GC content of 50.54% and Codon Adaptation index of 0.99 (Fig. 9). GC values fluctuating from 30 to 70 % are optimal and indicative of high gene expression in the host. Moreover, *E. coli* pET30a (+) vector was used for integration and optimal expression of the designed construct. For successful integration and cloning of the sequence into the vector, two restriction sites *Xho I* and *Nde I* were added with the help of SnapGene software (Fig. 10).

**Figure 9.**
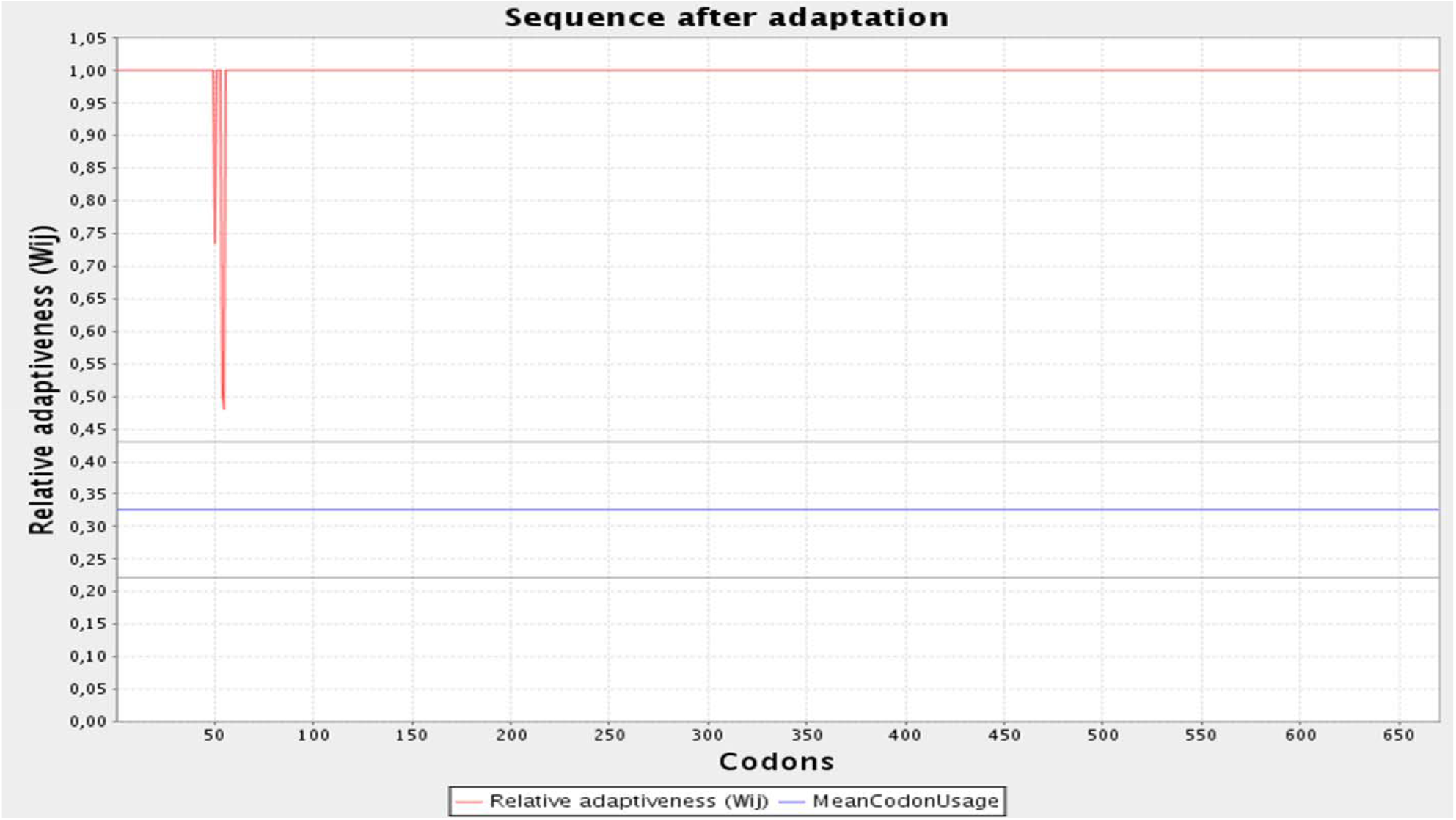
Adapted codons of the final vaccine protein. Codon adaptation index of optimized codon is 0.99 and GC value is found out to be 50.54%.

**Figure 10.**
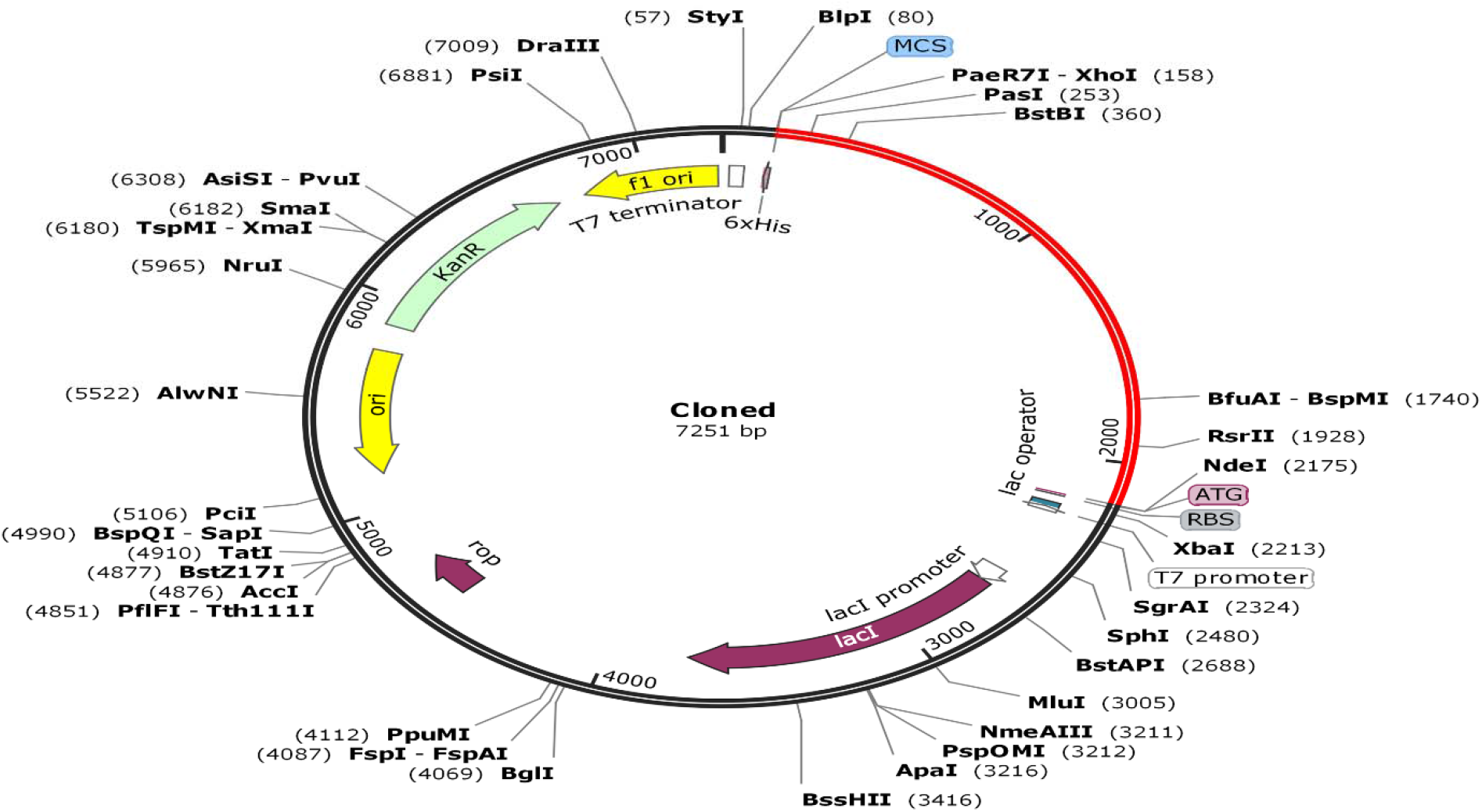
*In silico* restriction cloning of final protein into pET30a (+) vector. Red fragment signifies the inserted gene sequence of the proposed vaccine whereas black segment specifies backbone of E. coli vector. Cloned sequence is bounded by restriction enzymes *Xho I* and *Nde I*. 6x-His tag is positioned at C-terminus.

#### 3.3.11 Estimation of the Immune Responses

The collective immune responses after three times exposure to the antigen, produced by the final vaccine construct indicate a considerable spike in the levels of IgM as well as B-cell populations. Furthermore, an extortionate gush in the levels of IgM+ and IgG, IgG1, and IgG2 was also observed (Figure 11a). Recurrent exposures to the vaccine helped in the development of memory cells. Along with strong memory development, helper T cells and cytotoxic T cell populations showed decent levels of the increment (Figure 11 b, c, d).

**Figure 11.**
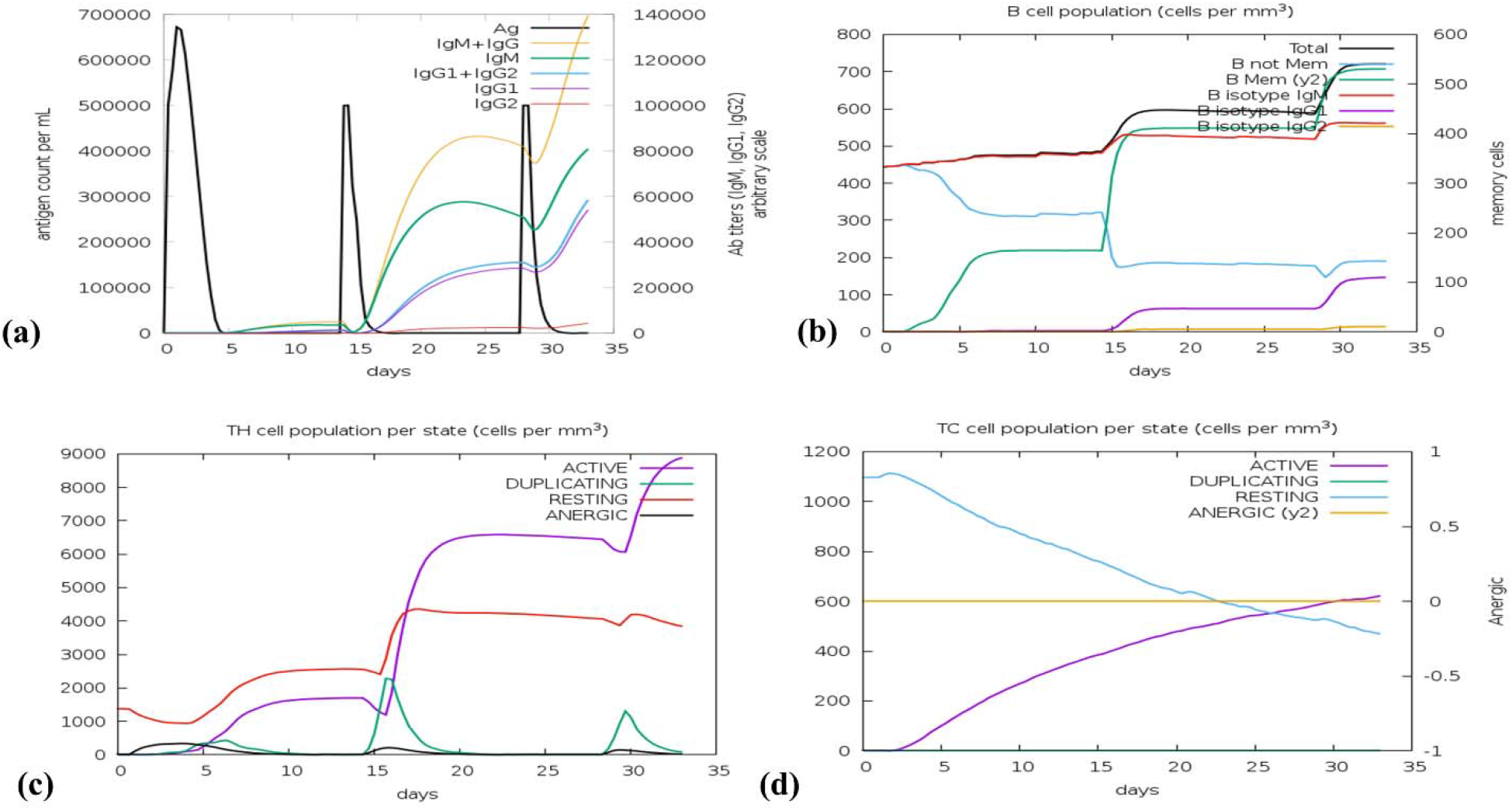
Simulated immune responses instigated by chimeric protein (a) Antibodies generated because of successive antigen injections; other immune cells are denoted by colored peaks (b) considerable shifts were observed in population levels of B-cells after three injections (c) Increments of helper-T lymphocytes (d) Cytotoxic T-cell populations per state after the injections. Cells that are not presented with the antigen are shown by the resting state while T-cells that demonstrate tolerance to the antigen because of repeated exposures are indicated by the anergic state.

## 4. Discussion

*A. baumannii* has emerged as an extremely cumbersome pathogen across the world becoming the leading cause of nosocomial and neonatal deaths (Peleg et al., 2008). It has successfully developed pan-drug resistance and thus appeared as one of the most difficult pathogens to treat or control (Styles et al., 2020). Almost 30 and 76% percent of the deaths are due to *A.baumannii* infections which can allegedly increase depending upon the severity of the patients (Ballouz et al., 2017). Different types of potential vaccine candidate have been discovered and many subunit vaccines consisting of protein or set of proteins have been studied against *A. baumannii* that include Outer Membrane Proteins (OMPs), Outer membrane Vesicles (OMVs), Inactivated Whole Cells Vaccines (IWCs), Ata, bap and several other subunit vaccines (Singh et al., 2016). However, due to the complex configurations, enhanced toxicity, and evocation of non-specific immune response, none of these vaccines could make their way to market or even the clinical trials (Ni et al., 2017a). More apt and rational vaccine design to combat this pathogen is the need of the hour. Therefore, we designed a multi-epitope vaccine for the first time using innocuous and more immunogenic regions of the outer membrane porin OprD.

The goal of this study was to introduce a novel multi-epitope vaccine design by exploiting numerous in-silico approaches against *A. baumannii*. OprD is a vaccine candidate protein predicted by the in-silico analysis of complete genomes and several strains of MDR *A. baumannii*. Complex proteome analysis of complete genomes and antibiotic-resistant strains of *A. baumannii* was performed by (Ni et al., 2017b). Outer membrane porins being profound in nature could instigate a considerable amount of immune response and induce immunization when encountered with *A. baumannii*. Eight porin proteins having potential roles as protective antigens and virulence factors were recognized, out of which one was OprD. Several studies support the prospective role of OprD as a successful vaccine candidate (Li et al., 2014). This porin participates in the transport of basic amino acids and carbapenem uptake. It is pertinent to use bioinformatics tools for carefully choosing the regions of OprD that confer immunogenicity. OprD is present in a wide array of clinical strains of *A.baumannii* (Ni et al., 2017a; Zhu et al., 2019). Using this protein would help in effectively targeting the complete range of pathogenic strains of *A. baumannii*.

This study focuses on generating a multi-epitope vaccine from a single representative protein of OprD. Latest computational tools for the development of vaccines have made it convenient to carefully select rationally apt epitopes from the proteomes of pathogenic microbes. Employment of such tools could help us pave ways towards designing an effective vaccine strategy. To have a reliable vaccine model, it is germane to bring into account all the facets of vaccine development viz. secondary and tertiary structure, antigenicity, allergenicity, flexibility, accessibility, and hydrophilicity of the vaccine. Sticking to only one of these characteristics would not generate the needed results.

The antibody-mediated or the humoral immune response is prompted by the recognition of distinct features of the pathogen considered as non-self by the host (Chaplin, 2010). These distinct features are antigenic determinants, also known as epitopes that are specifically detected by the antigen-binding site of the antibodies present in the host’s immune system (Cruse and Lewis, 1999). There have been studies supporting the role of B cells and T cells immunity against *A.baumannii* infections (Chen, 2020; Morris et al., 2019). The role of the TLR4 pathway has also been reported in detail along with the potential roles of other TLRs in mediating the interaction between antigens and the host immune system (Kim et al., 2013). We first projected T cell and B cell epitopes of the protein and combined them with the aid of specific linkers to obtain a multi-epitope vaccine as a result. These spacers play a key role in improving vaccine design (Shey et al., 2019). Formerly reported cleavable linkers, AAY, GPGPG, and EAAK were fused into the final vaccine design with AAY and GPGPG connecting the forecasted peptides and EAAK linking the adjuvant to the N terminal of the B cell epitopes (Dong et al., 2020). A six-His tag was joined at the C-terminus to ensure purification at the later stages. Several computations based on immunoinformatic analysis revealed that the designed vaccine model contains high-affinity MHC class I and class II epitopes and linear B cell epitopes in bulk amounts. The absence of allergenic features in this model further endorses the safety of it as a vaccine candidate.

The molecular weight of our construct was observed to be 69.9 kDa with the capability of being soluble when expressed. Apart from defining the bioactivity of the protein, the solubility of protein also plays a crucial role in the biochemical and functional analysis (Ahmad et al., 2018). The protein was detected to be slightly acidic with a theoretical pI of 5.51. Similarly, it was noted that the protein will be stable upon expression, hence reinforcing the competence of the designed construct. The protein was also observed to be hydrophobic due to the presence of aliphatic side chains as indicated by the aliphatic index value. All these features designate the protein as has ability to withstand high temperatures and thus well suited for utilization in the endemic parts.

Information regarding the secondary, as well as the tertiary structure, is vital in vaccine designing (Shey et al., 2019). Secondary structure analysis revealed that the protein is largely comprised of coils (43%), with only 7.16% of the residues disordered. The 3D structure of the vaccine showed notable refinement and thus resulted in the attainment of the desirable properties such as Ramachandran plot values. Rama-favored regions had a score of 94.5 % with rare residues in the outlier region thus suggesting the high quality of the model.

Energy minimization was performed to stabilize the overall conformation as well as minimize the potential energy of vaccine. During energy minimization, anomalous parts of the structure are repaired hence giving rise to a more stable protein structure that would behave more efficiently in the life-like cellular environments. The predicted RMSD of the vaccine-TLR-4 complex was predicted to be 0.5Å, which confirms the firmness of the complex.

Several studies have demonstrated the role of TLR4 in protecting against *A.baumannii* infections. Therefore, a data-driven docking analysis was carried out to assess the possible interactions between the designed construct and TLR4. Results suggest that the designed construct could effectively stimulate the immune response. Binding energies elucidate the strong binding of TLR4 with the construct. It is strongly recommended to further investigate the interactions between TLR4 and the construct in vitro as well as in vivo.

The results obtained by immune simulations complied with the actual immune responses. A continuous spike in immune responses was observed after giving repeated exposures to the antigen.IgG1, IgG3, IgE immune responses have been reported against *A.baumannii* in different reports (Ansari et al., 2019; Cha et al., 2018). We observed a spike in B and T cell populations, as well as a considerable amount of peak in Th cell populations, was also noted.

The first and foremost step for the validation of a potential vaccine candidate is to inspect it for immunoreactivity (Gori et al., 2009). For this purpose, it is necessary to express the target protein in an appropriate host. *E. coli* is a renowned expression system to attain maximum protein expression levels (Chen, 2012). To achieve high-level expression in *E.coli* (strain K12), codon optimization was executed. The values of the CAI (0.99) and GC content (50.54%) were well-suited to obtain an optimum expression.

## 5. Conclusion

Novel and effective strategies are required to cope with difficult-to-treat *A. baumannii* infections. This study makes use of several in-silico tools to design a rational vaccine that consists of multiple B and T cell (CTL and HTL) epitopes. Our designed model is more antigenic and immunogenic with decreased cytotoxic effects which could trigger antibodies associated protection.

## 6. Disclosure statement

No conflict of interest was reported by the authors.

## Acknowledgment

The authors would like to acknowledge the computational help to run molecular dynamics simulations by Lab Engineer Muhammad Hassan Khan from Research Center for Modeling and Simulation (RCMS), National University of Sciences and Technology (NUST), Islamabad, Pakistan.

## Notes

### Competing Interest Statement

The authors have declared no competing interest.

